# Model-Based Recommendations for Optimal Surgical Placement of Epiretinal Implants

**DOI:** 10.1101/743484

**Authors:** Michael Beyeler, Geoffrey M. Boynton, Ione Fine, Ariel Rokem

## Abstract

A major limitation of current electronic retinal implants is that in addition to stimulating the intended retinal ganglion cells, they also stimulate passing axon fibers, producing perceptual ‘streaks’ that limit the quality of the generated visual experience. Recent evidence suggests a dependence between the shape of the elicited visual percept and the retinal location of the stimulating electrode. However, this knowledge has yet to be incorporated into the surgical placement of retinal implants. Here we systematically explored the space of possible implant configurations to make recommendations for optimal intraocular positioning of the electrode array. Using a psychophysically validated computational model, we demonstrate that better implant placement has the potential to reduce the spatial extent of axonal activation in existing implant users by up to ∼55 %. Importantly, the best implant location, as inferred from a population of simulated virtual patients, is both surgically feasible and is relatively stable across individuals. This study is a first step towards the use of computer simulations in patient-specific planning of retinal implant surgery.

## 1 Introduction

Argus II (Second Sight Medical Products, Inc., https://secondsight.com) is currently the only retinal prosthesis system to receive approval from both the US Food & Drug Administration and the Conformité Européenne Mark. For successful implantation of the device, surgeons are instructed to place the electrode array parafoveally over the macula, approximately diagonal at −45° to the horizontal meridian (see Surgeon Manual [10], p.29), for reasons of surgical ease.

However, instead of seeing focal spots of light, patients implanted with epiretinal electronic implants report seeing highly distorted percepts that range in description from ‘blobs’ to ‘streaks’ and ‘wedges’ [8]. Electrophysiological evidence from *in vitro* preparations of rat and rabbit retina suggests that these distortions may arise from incidental stimulation of passing axon fibers in the optic fiber layer (OFL) [5,6,11]. Although there is believed to be a systematic relationship between the severity of distortions due to axonal stimulation and the retinal location of the stimulating electrode [2,3,4], this knowledge has yet to be incorporated into the patient-specific planning of retinal implant surgery [1], or surgical recommendations for intraocular positioning of the electrode array.

The contributions of this paper are three-fold. First, we present a strategy to optimize the intraocular placement of epiretinal implants, based on a psychophysically validated computational model of the vision provided by Argus II. Second, we validate this strategy on three Argus II patients and a population of virtual patients. Third, we recommend an optimal intraocular location that is both surgically feasible and relatively consistent across individuals.

## 2 Methods

Prior work suggests a dependence between the shape of a visual percept generated by an epiretinal implant and the retinal location of the stimulating electrode [4,5]. Because retinal ganglion cells (RGCs) send their axons on highly stereotyped pathways to the optic nerve [7], an electrode that stimulates nearby axonal fibers would be expected to antidromically activate RGC bodies located peripheral to the point of stimulation, leading to percepts that appear elongated in the direction of the underlying nerve fiber bundle (NFB) trajectory (Fig. 1, *right*) [4]. As can be seen in (Fig. 1, *left*), electrodes near the horizontal meridian are predicted to elicit circular percepts, while other electrodes are predicted to produce elongated percepts that will differ in angle based on whether they fall above or below the horizontal meridian.

**Fig. 1.**
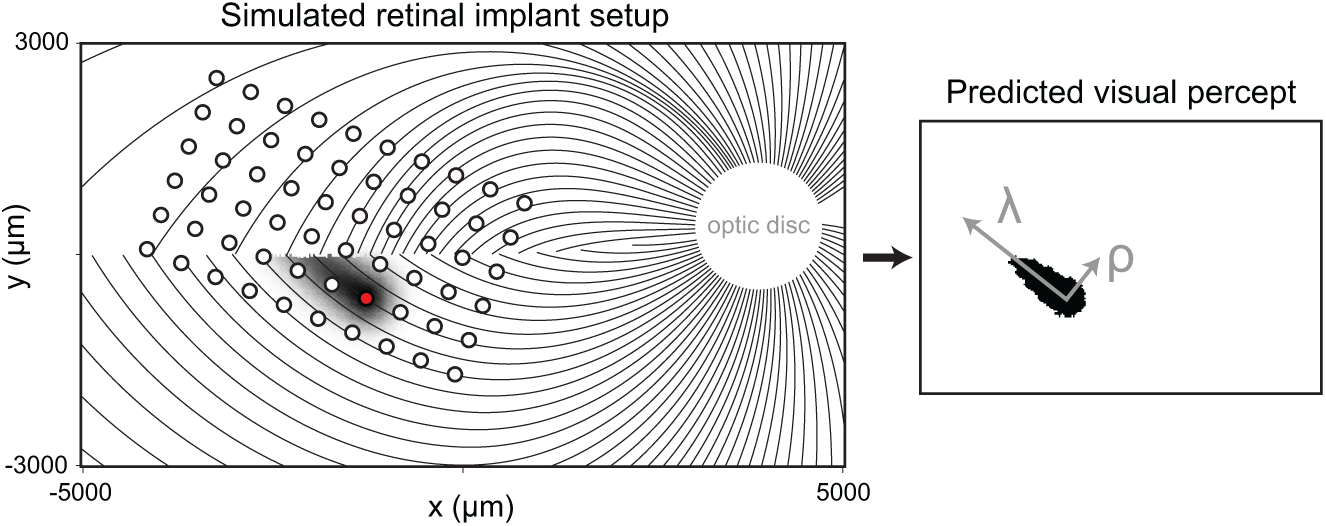
A simulated map of retinal NFBs (*left*) can account for visual percepts (*right*) elicited by epiretinal implants. *Left*: Electrical stimulation (red circle) of a NFB (black lines) could antidromically activate retinal ganglion cell bodies peripheral to the point of stimulation, leading to tissue activation (black shaded region) elongated along the NFB trajectory away from the optic disc (white circle). *Right*: The resulting visual percept appears elongated as well; its shape can be described by two parameters, *λ* (spatial extent along the NFB trajectory) and *ρ* (spatial extent perpendicular to the NFB). See [4] for more information.

Ref. [4] used a simulated map of NFBs in each subject’s retina to accurately predict the shape of percepts elicited by the Argus system, assuming that:

i. An axon’s sensitivity to electrical stimulation decays exponentially with decay constant *ρ* as a function of distance from the stimulation site (*x*_stim_, *y*_stim_).
ii. An axon’s sensitivity to electrical stimulation decays exponentially with decay constant *λ* as a function of distance from the soma (*x*_soma_, *y*_soma_), measured as path length along the axon.

This allowed for percept shape to be described as a 2-D intensity profile, *I*(*x, y*):

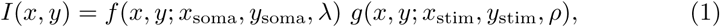

where *f* modeled an exponential fall-off along the axon, with maximal sensitivity close to the cell body:

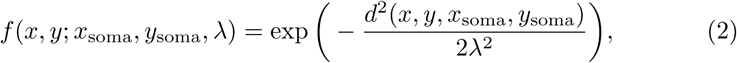

(using path length *d*(*x, y, x*_soma_, *y*_soma_) measured between a point (*x, y*) on the axon and the soma (*x*_soma_, *y*_soma_); and *g* was a two-dimensional Gaussian function centered over (*µ, ν*) with standard deviation *σ*:

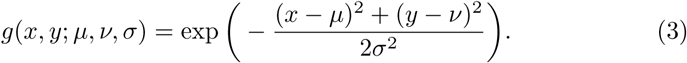

The resulting intensity profile was then thresholded to arrive at a binary image, which served as the predicted visual percept (Fig. 1, *right*).

This model was previously validated on psychophysical data from three Argus II patients with severe retinitis pigmentosa [4]. Electrical stimulation was delivered to a number of pre-selected electrodes in random order, and subjects were asked to outline perceived percept shape on a touch screen. The images predicted by the model were then compared to the drawings, and the best-fitting values for *ρ* and *λ* were determined for each subject in a cross-validation procedure. Note that a single value of *ρ* and *λ* was fitted for each subject, and then used for all electrodes in that subject’s array (see Table 1).

**Table 1.** Model parameters. Device placement and optic disc location were estimated from fundus photographs, whereas *ρ* and *λ* were fit to psychophysical data (see [4] for details). Device rotation was measured with respect to the horizontal meridian (positive angles: counter-clockwise rotation). The fovea was located at (0, 0).

| Subject ID | Device center<br>($x, y$; $\mu\text{m}$) | Device rotation<br>(deg) | Optic disc center<br>($x, y$; deg) | $\rho$<br>($\mu\text{m}$) | $\lambda$<br>($\mu\text{m}$) |
| --- | --- | --- | --- | --- | --- |
| 1 | (−1331, −850) | −28 | (16.2, 1.4) | 352 | 299 |
| 2 | (−467, 206) | −26 | (14.0, 1.2) | 91 | 659 |
| 3 | (−1807, 401) | −22 | (16.3, 2.4) | 414 | 1383 |

To determine the optimal intraocular positioning of Argus II for Patients 1–3, we performed a grid search over the space of feasible implant configurations and used the model described in [4] to estimate average percept size. We limited the search to a region of the retina where the model was deemed valid [7]. This included array centers located in the range *x* ∈ [− 2000 µm, 400 µm] and *y* ∈ [− 1200 µm, 1200 µm], which we sampled at 200 µm resolution. We considered implantation angles in the range [− 90°, 90°] with a 5° step size.

Since *ρ* and *λ* were fixed for each subject, the size of each predicted percept was closely related to the amount of axonal stimulation. Moreover, since visual outcomes in epiretinal implants depend crucially on the ability of the device to generate localized percepts, average percept size serves as a simple proxy for the quality of the generated visual experience.

## 3 Experiments and Results

### 3.1 Optimal Implant Placement: Three Argus II Patients

Results are shown in Fig. 2. Percept predictions for the actual implant configuration are shown in the leftmost column, where each percept is overlaid over the corresponding electrode in a schematic of each patient’s implant.

**Fig. 2.**
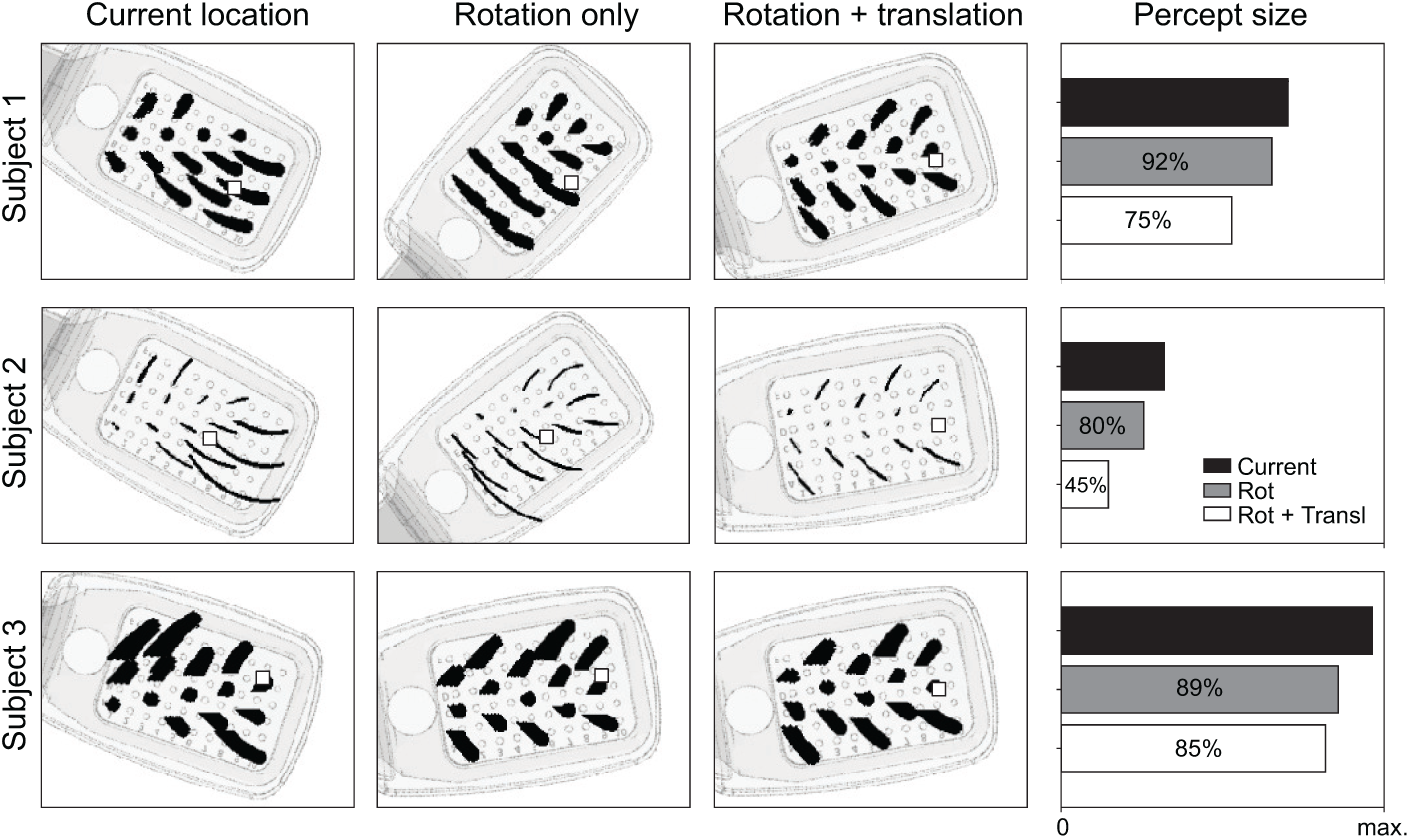
Model predictions of percept shape for different electrodes, overlaid over a schematic of each patient’s implant. Predictions for the actual implant configurations (leftmost column) are contrasted with optimized arrangements, either where the device location is the same but device orientation is adjusted to minimize the spatial extent of axonal activation (second column), or where both device location and orientation are optimized (third column). Mean percept size for the three configurations are shown in the rightmost column. Small squares indicate the location of the fovea.

Consistent with the psychophysical data described in [4], electrodes located in close proximity to the horizontal meridian elicited more focal percepts than more eccentric electrodes. One could therefore reduce average percept size without changing the location, simply by rotating the array so that as many electrodes as possible lie close to the horizontal meridian (second column, labeled “rotation only”); a strategy that worked especially well for Patient 2.

On the other hand, if one were free to place the implant at any parafoveal location oriented at any angle in [−90°, 90°], average percept size could be even further reduced (third column, labeled “rotation + translation”).

The possible reduction in percept size is quantified in the rightmost column. In the case where the implant location was fixed, but a rotation of the device was allowed, mean percept size could be reduced by up to 20 %. When both location and angle were allowed to vary the mean percept size could be reduced by up to 55 %. Interestingly, Fig. 2 suggests that all three patients could have benefited from a similar intraocular positioning of the device, a roughly 90° shift from the currently recommendation location.

Fig. 3 further quantifies the effect of array positioning on mean percept size. Heat maps in the left column show the effect of altering the array location. At each location, the implant was rotated to find the angle that minimized average percept size. These corresponding rotation angles are shown in the right column. Thus, for Patient 1, the ideal location would be *x* = − 2000, *y* = 0, at a rotation of ∼20°. For all three patients, the ideal location lay close to *x* = − 2000, *y* = 0. Indeed, by happenstance, Patient 3’s device lay close to the optimal implant location. However, due to specifications by the device manufacturer, all three patients had the array implanted at negative angles almost 45° away from the optimal angle. Our simulations suggest that positive angles (reddish colors) would be preferable for most implant locations.

**Fig. 3.**
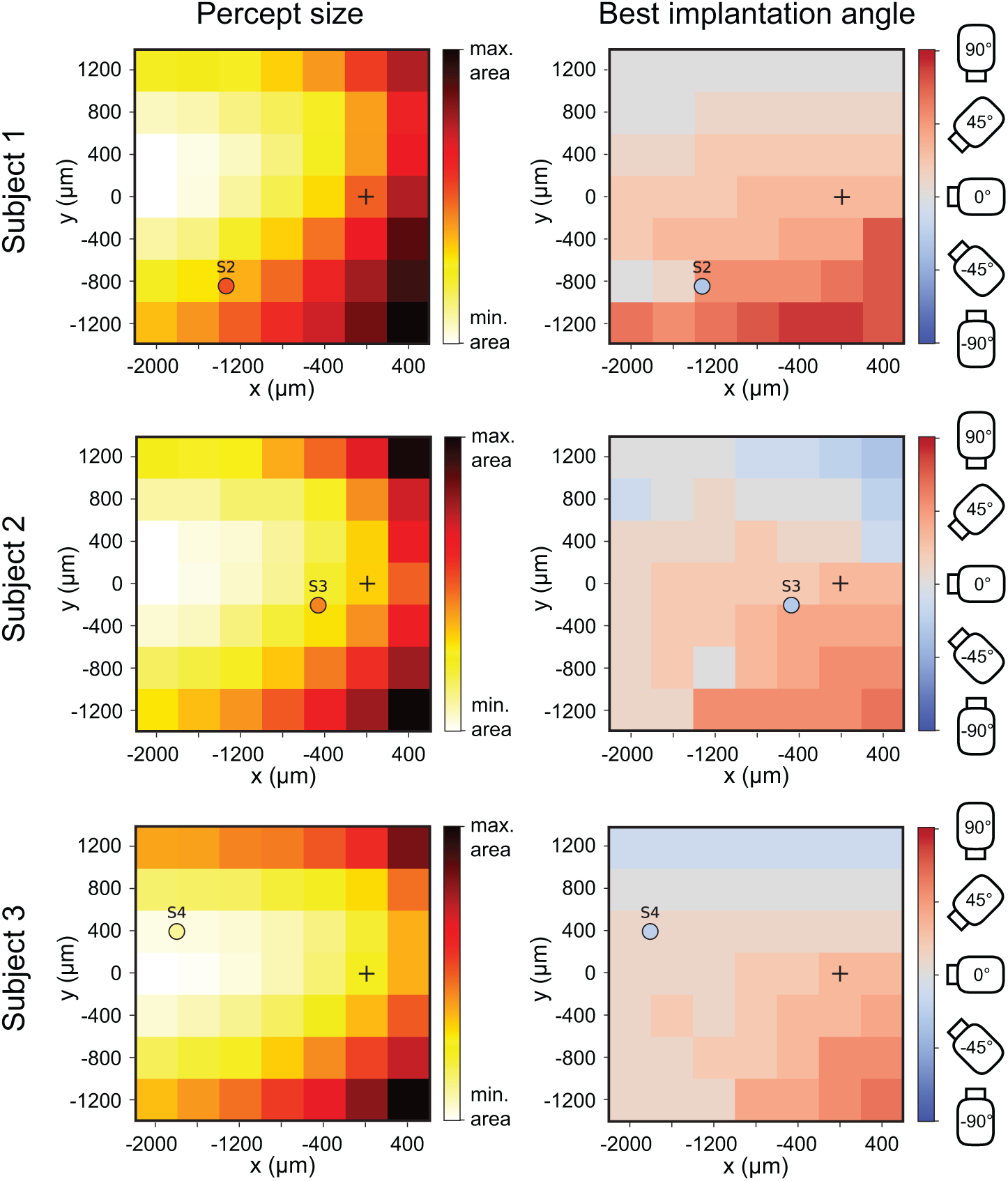
Percept size for different implant configurations for Patients 1–3. *Left*: Surfaces in each panel show the minimal percept size that could be achieved by optimal device rotation, as a function of device location (i.e., center of the array). Small circles depict actual device locations for Patients 1–3, with the color indicating the empirically measured percept size. *Right*: The corresponding device rotation angle used to achieve minimal percept size in the left column, with the color of the small circles indicating the actual device rotation for Patients 1–3. Note that due to the symmetry of the electrode grid in Argus II, 90° and *-*90° are two technically equivalent configurations. The small black cross indicates the fovea.

### 3.2 Optimal Implant Placement: Virtual Patients

To investigate whether these findings would generalize to other Argus II users, we simulated a population of 90 virtual patients, each with randomly assigned values for *ρ* (in the range [50 µm, 500 µm]) and *λ* (in the range [200 µm, 1600 µm]). These two model parameters are thought to capture important individual differences across patients, and might serve as a phenomenological description of both neuroanatomical parameters as well as drawing preferences [4]. We then repeated the experiment described above to determine the optimal implant location and orientation for each individual in the population.

The result is shown in Fig. 4. Here, each value in the heat map is the median value obtained across all virtual patients for that particular implant location. Interestingly, the findings described in Fig. 2 above also hold for the entire population of virtual patients, suggesting that it is not important to know *ρ* and *λ a priori*. This is crucial, as the values for *ρ* and *λ* can currently only be measured after successful implantation using psychophysical paradigms.

**Fig. 4.**
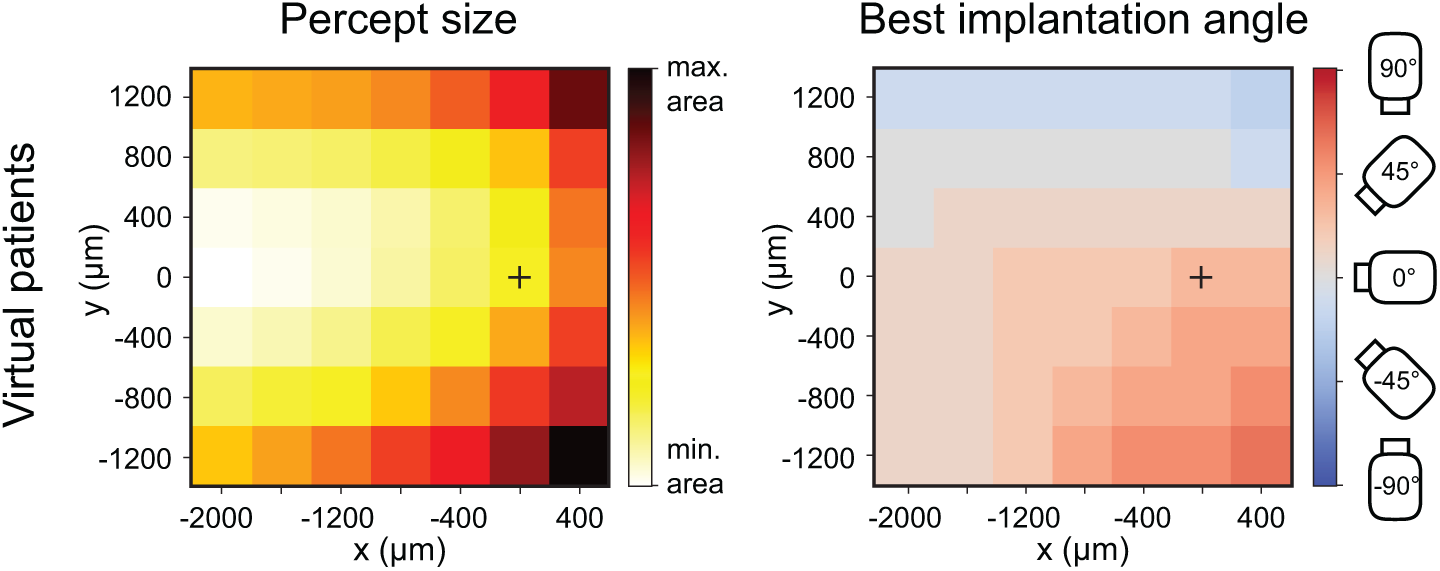
Percept size for different implant configurations in a population of virtual patients. *Left*: Surfaces in each panel show the minimal percept size that could be achieved by optimal device rotation, as a function of device location (parameterized by the location of the center of the array). *Right*: The device rotation angle used to achieve minimal percept size in the left column. The small black cross indicates the fovea.

These simulations indicate that an epiretinal implant should best be placed at ∼2000 µm temporal to the fovea, centered over the horizontal meridian, and orientated at ∼10° Importantly, this configuration is surgically feasible, and in fact has been achieved before [1]. On the one hand, optimal implantation angle changes only gradually with location (Fig. 4, *right*), underscoring the practical feasibility of this approach. On the other hand, percept size is more sensitive to implantation angle in the optimal location than elsewhere (full width at half maximum in optimal location: ∼60°, elsewhere: ∼90°), underscoring the importance of choosing a suitable implantation angle.

## 4 Conclusion

This preliminary study is a first step towards the use of computer simulations in the patient-specific planning of retinal implant surgery. We show here that the visual outcome of epiretinal implant surgery might be substantially improved by guiding the intraocular positioning of the electrode array using a patient-specific computational model of the spatial layout of the OFL. Our findings suggest that optimized array placement could reduce the spatial extent of axonal activation in existing Argus II users by up to ∼ 55 %. Importantly, predicted percept sizes are robust to small deviations from the optimal location and orientation of the array, as well as to small deviations of the model parameters that predict the shape of the percept.

The optimal implant location, as inferred from a population of virtual patients, is ∼2000 µm temporal to the fovea, centered over the horizontal meridian, and orientated at ∼10° with respect to the meridian. Importantly, this placement is surgically feasible. Our method requires *a priori* knowledge about the location of the fovea and horizontal meridian, but these can be estimated presurgically based on anatomical landmarks in fundus images. Moreoever, intraoperative optical coherence tomography (OCT) can be used to guide the placement of the array [9].

## Acknowledgments

Supported by the Washington Research Foundation Funds for Innovation in Neuroengineering and Data-Intensive Discovery (M.B.), by an award from the Gordon and Betty Moore Foundation (Award #2013-10-29) and the Alfred P. Sloan Foundation (Award #3835) to the University of Washington eScience Institute (A.R.), and by the National Institutes of Health (NIH K99 EY-029329 to M.B., EY-12925 to G.M.B., and EY-014645 to I.F.). Research credits for cloud computing were provided by Amazon Web Services.

